# Towards A 3D Chromosome Shape Alphabet

**DOI:** 10.1101/2020.08.04.236224

**Authors:** Carlos Soto, Darshan Bryner, Nicola Neretti, Anuj Srivastava

## Abstract

The study of the 3-dimensional (3D) structure of chromosomes – the largest macromolecules in biology – is one of the most challenging to date in structural biology. Here, we develop a novel representation of chromosomes, as sequences of *shape letters* from a finite *shape alphabet*, which provides a compact and efficient way to analyze ensembles of chromosome shape data, akin to the analysis of texts in a language by using letters. We construct a *Chromosome Shape Alphabet* (CSA) from an ensemble of chromosome 3D structures inferred from Hi-C data – via SIMBA3D or other methods – by segmenting curves based on topologically associating domains (TADs) boundaries, and by clustering all TADs’ 3D structures into groups of similar shapes. The median shapes of these groups, with some pruning and processing, form the *Chromosome Shape Letters* (CSLs) of the alphabet. We provide a proof-of-concept for these CSLs by reconstructing independent test curves using only CSLs (and corresponding transformations) and comparing these reconstructions with the original curves. Finally, we demonstrate how CSLs can be used to summarize the variability of shapes in an ensemble of chromosome 3D structures using generalized sequence logos.

## 1 Introduction

Among the most complex structural problems in biology is the study of how the mammalian genome, measuring approximately 2 m, is organized within a volume of roughly 10^−16^ m^3^, the cell nucleus. New techniques developed over the past 10-15 years have allowed us to take a closer look at the genome’s 3D organization. By studying contact matrices generated by proximity ligation assays such as Hi-C [7], we now know that each section of the genome belongs to one of two main compartments, called A and B, which are comprised of several sub-compartments [15]. Genomic regions within the same sub-compartment tend to interact with each other and avoid interactions with other sub-compartments. Locally, chromosomes are folded into regions of highly interacting loci, referred to as Topologically Associating Domains (TADs), which are thought to define units containing genes and their regulatory elements [21]. We can also use the Hi-C contact matrices to infer the 3D structure of chromosomes and ultimately of the whole genome [12]. Recent studies, including single cell Hi-C [9, 10] and microscopy [11, 1, 8], have demonstrated that 3D structural variability exists not only across cell types but also within the same cell type, raising the question: *What structural features define the phenotype of a cell*? However, due to the high dimensionality of the state space of 3D curves, this is a very hard problem to tackle, and new methodologies are needed to cast this problem into a more manageable framework. To address this problem, we develop a new representation of chromosomes structures based on the following hypothesis on their variability:

### Hypothesis

*Most 3D chromosome structures can be obtained by concatenating a limited number of basic structural units, termed* **Chromosome Shape Letters** *or CSLs. Different chromosomes use the same CSLs but differ in frequencies and placements of these letters in their organizations across cells, much like sentences in a language*.

We shall call a set of CSLs a Chromosome Shape Alphabet (CSA). Our aim in this paper is to demonstrate how to construct such an alphabet and to demonstrate its applications in characterizing chromosome populations.

The main motivation of this hypothesis is as follows. Although chromosomes structures exhibit tremendous variability across species, cells, and strains, they share certain common substructures at local, smaller scales. For example, all chromosomes contain certain spiral structures – single loops, double loops, etc. – and some simple connecting threads that bind these intricate structures. This commonality of substructures is striking both visually and quantifiably in estimated 3D structures, irrespective of the package or methodology used in estimating chromosome structures. Also, while global genomic and chromosome structures are vastly complex and highly variable, the structural units are usually relatively simple and quite stable.

*What are the implications of this hypothesis*? If this hypothesis holds then a chromosome, and indeed the full genome, can be symbolically represented by a string of letters, each representing a shape letter, in a manner that is similar to forming words or sentences from a chosen alphabet. (One would also need a set of transformation variables – rotation, scale, etc. – that govern concatenations of CSLs into full 3D curves.) The consequences of such a representation are enormous. This can be a very useful tool in coding structural genomic variability and analyzing it in simplified, abstract terms. One can estimate chromosome structures from Hi-C or imaging data, directly in terms of their CSLs, much like the estimation of letters or words in speech signal processing. Similar to explorations of vast linguistic variability, exhibited in written and spoken media, using hierarchical organizations of a very few letters (only 26 for the English language), one can develop a very comprehensive toolset for exploring complex genomic structural universe. For instance, one can study relative frequencies and co-frequencies of these CSLs in a chromosome population. Or, use these frequencies in studying effects of genetic mutations and other biological events on genomic structures.

This is a novel research direction with little precedence in the current biology literature. There are some broad structural labels associated with secondary structures of proteins, e.g. *α* helices, *β* sheets, turns, and loops. There also exists the notion of building anatomical atlases for analyzing commonality and variability of structures in medical imaging literature. However, there are currently no such structural labels or atlases for chromosomes. Furthermore, we are aiming for a detailed, comprehensive listing of substructures that can used to completely characterize full 3D chromosomes. For instance, we would like to reconstruct full chromosomes and even full genomes using these shape letters and associated transformations. These aspects separate the current paper from past shape analyses in bioinformatics or biology literature. The proposed framework can also be applied to represent analyze other objects such as other polymers, tertiary structures of a protein, or RNA.

The next question is: *How to investigate and validate this hypothesis involving chromosome shape letters*? Such an investigation requires a whole suit of concepts, some of them being very novel to the field. The development of these concepts listed below becomes the main contribution of this paper.

- **Resolution-Specific Definition of CSA**: Firstly, one requires a proper scientific definition of CSLs for chromosomes of interest. To break a chromosome into its structure units depends on the chosen resolution, *i.e*. one can chop a chromosome into a few or many units. We will use the notion of topologically associated domains, or TADs, to specify the resolution of at which CSAs are defined. TADs provide segmentations of full chromosomes into their smaller units and that will form a basis for our definition of CSA. Since TAD definitions are not universal, the subsequent definition of a CSA is potentially dependent on how TADs are defined. The specification of resolution also relates to the resolution at which the underlying contact matrices are constructed.
- **Data-Driven Discovery of CSLs**: One can either derive CSLs from a theoretical perspective, using purely energy-based, biological arguments, or use an empirical approach, exploring vast amount of previously estimated 3D conformations for common substructures. Taking the latter approach, we shall explore and collate substructures common in training data and use them to define our CSLs.
- **Complexity-Based Nomenclature**: Once a number of potential shapes have been collected, we need to organize and label them mnemonically for use as letters. We use a complexity-based ordering of collected shapes, pruned appropriately to avoid redundancy, and give them letter names – `A, B, C`, etc. Analogous to a language, we can then use these letters to represent and analyze chromosome shapes.
- **CSA Representation**: One needs tools to represent individual chromosomes as sequences of CSLs and transformations. In other words, given a test curve 3D chromosome curve, one needs to select which CSLs, in what order, and under which transformations, can reconstruct that original chromosome. Further, this procedure allows us to quantify reconstruction errors and to evaluate the performance of a CSA in representing future 3D chromosome data.
- **CSA Validation**: There are several perspectives for validating such atomic representations of biological structures. One can do it from a *biological* perspective by providing biological interpretations to these representations and using them to address pertinent biological questions. The other approach is purely *geometric*, where one compares original and reconstructed structures in terms of their shapes or some other objective function. In this paper we will follow a geometric perspective and leave the biological perspective as future research.

In order to organize and refine a set of segmented chromosomes into a CSA, we will need tools from shape analysis of 3D curves. For this there are several choices, and we will use *elastic shape analysis* as described in [20]. This approach compares shapes of curves using a proper metric and allows for computing statistical summaries of curves. An important advantage of this approach is that it does not presume registration of points across curves, but finds it during shape analysis. This is important in the current study because not all chromosomes are chopped into TADs in a precise, synchronized fashion, and one needs to register them geometrically for shape comparisons. We need a large number of native chromosome conformations to perform this data-driven discovery of CSLs. In this paper we utilize the SIMBA3D package [16] for estimating chromosome structures from Hi-C data. In recent years, Hi-C [12] has emerged as an important sensing modality for learning chromosome structures. Hi-C data can be collected from either a single cell or an ensemble of cells. Accordingly, a number of methods have been developed to estimate chromosome structures from the Hi-C contact data. These methods differ in their problem formulations, the choices of objective functions and the techniques used for optimizations. Examples of statistical solutions for chromosome structure estimation include MCMC5C [17], BACH [6], SIMBA3D [16], and tRex [13]. Although we use SIMBA3D here, any of these other packages can easily be substituted in the process.

The rest of this paper is laid out at follows. We start by providing a brief summary of relevant tools from elastic shape analysis of 3D curves in Section 2. Then we lay out steps of our data-driven discovery of CSAs using past 3D chromosome data in Section 3. These steps include segmenting full curves using TADs, clustering and summarizing shapes of these segmented curves, and building a collection of distinct shape letters. Section 4 presents a validation of the underlying hypothesis in curve reconstruction, *i*·*e*. reconstructing curves using only the CSLs. Section 5 illustrates some uses of a CSA in characterizing a chromosome population. The paper ends with a short discussion in Section 6.

## 2 Elastic Shape Analysis Tools

In this section we summarize a set of useful tools from elastic shape analysis of 3D curves. These tools are used to compare, cluster, summarize, and collate curves according to their shapes. We will apply these tools to segments obtained by chopping 3D curves representing chromosome conformations.

### 2.1 Shape Representation and Metric

Let ℱ represent the set of all smooth maps from [0, 1] to ℝ^3^. An *f* ∈ ℱ represents a parameterized 3D curve. Given arbitrary elements of ℱ, our goal is to compare and analyze their shapes. Shape is a property that is invariant to *sizes, rigid motions and parameterizations* of these curves. In other words, if we scale, rotate, translate or re-parameterize a curve, its mathematical representation changes but its shape remains unchanged. Let 𝕆 (3) denote the set of all 3 *×* 3 orthogonal matrices – it is the set of all rotations and reflections in 3D. For any *O* ∈ 𝕆 (3), the curve *O · f* denotes a rotation/reflection of the original curve *f* ∈ ℱ. Also, let Γ = {*γ* : [0, 1] → [0, 1]|*γ*(0) = 0, *γ*(1) = 1, *γ* is a diffeomorphism}. For any *γ* ∈ Γ, the curve (*f* ∘ *γ*)(*t*) ≡ *f* (*γ*(*t*)) is called a re-parameterization of *f*. The shape of a curve *ρO · f* (*γ*(*t*)) + *x*, for any *ρ* ∈ ℝ_+_, *x* ∈ ℝ^3^, and *γ* ∈ Γ is exactly the same as that of *f*. Elastic shape analysis is a powerful framework developed to ensure this invariance in all computations. It defines a new representation called the *square-root velocity function* (SRVF) of a curves. For a curve *f* ∈ ℱ, its SRVF is given by 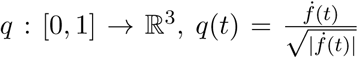, where 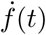 is the velocity vector at the point *f* (*t*). For a certain class of smooth functions, their SRVFs are elements of 𝕃^2^, the set of all square-integrable functions on [0, 1].

If the SRVF of a curve *f* is *q*, then the SRVF of the curve *O* · (*f* ∘ *γ*) is 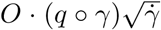. Also, one can check that the length of *f* is same as the 𝕃^2^ norm of *q*. So, if we want to rescale a curve *f* to be of unit length, we can simply rescale the corresponding SRVF according to *q* ⟼ *q/* ‖ *q* ‖. The set of all scaled SRVFs is an infinite-dimensional unit sphere 𝕊_∞_. The shape of a curve *f* is represented by the set of all possible rotations, reflections, and re-parameterizations of its SRVF: 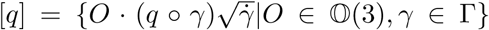. This is also called an *orbit* of *q* under rotations and re-parameterizations. The set of all orbits is called the *shape space S* = {[*q*]|*q* ∈ 𝕊_∞_}.

Given any two curves *f*_1_ and *f*_2_, with SRVFs *q*_1_ and *q*_2_, respectively, the difference in their shapes is given by:

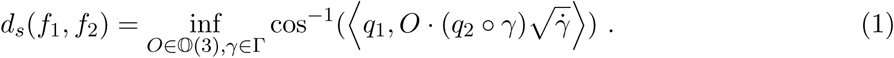

The infimum is computed numerically using the *Procrustes alignment* and the *Dynamic Programming Algorithm*. The quantity *d*_*s*_ is a proper shape metric in that sense that it satisfies all properties, including the triangle inequality, on the shape space *S*. For any two curves *f*_1_ and *f*_2_, one can also compute the shortest path between their shapes in *S*. These paths are called *geodesics* and provide a way to optimally deform one shape into another. Figure 1 shows some examples of geodesic deformations where in each row, the first curve is *f*_1_ and the last curve is *f*_2_, and the intermediate curves represent the geodesic path that deforms *f*_1_ into *f*_2_.

**Figure 1:**
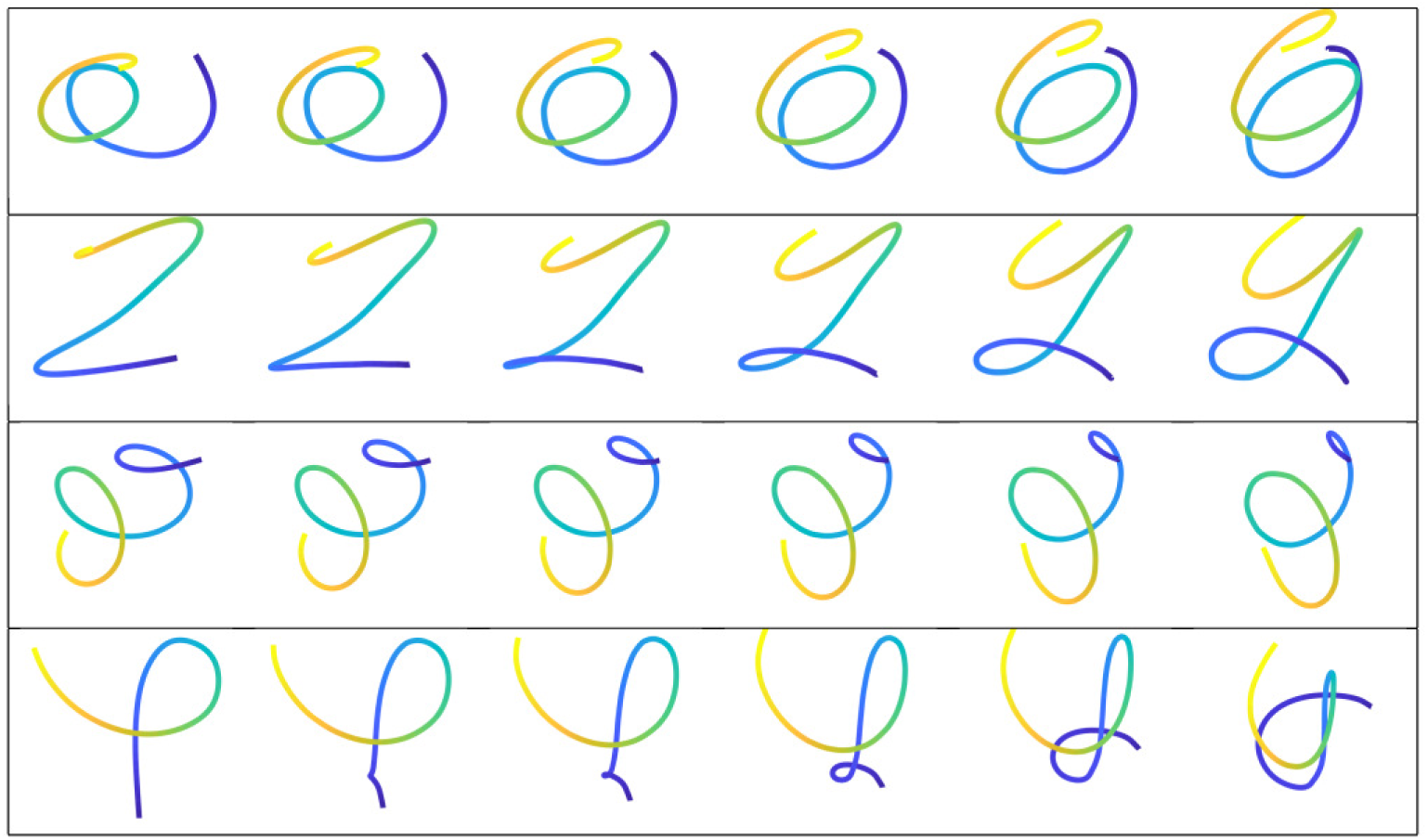
Examples of geodesic deformations between curves *f*_1_ (leftmost) and *f*_2_ (rightmost) in each row.

For readers that are not familiar with elastic shape analysis, we recommend the textbook [20] for a detailed treatment. It describes the motivation for using SRVF representation of curves and many of the nice mathematical and computational properties associated with SRVF representations.

One way this study differs from the classical shape analysis is that it includes reflections, in addition to the rotations, for comparing shapes of any two curves. In the classical shape analysis, one rotates a structure to best align with the other structure, while comparing their shapes. The reflections are not allowed. However, given the nature of the Hi-C data and the procedures used in estimating structures from Hi-C data, we extend the transformation space from rotations (𝕊 𝕆 (3)) to rotations and reflections (𝕆 (3)). We illustrate this point further using an example in Fig. 2. In the first two panels, we show two curves *f*_1_ and *f*_2_. In the third panel, we show the optimal rotational alignment of *f*_2_ to *f*_1_, *i.e*. search over 𝕊 𝕆 (3). In the last panel, we show the best rotational and reflectional alignment of *f*_2_ to *f*_1_, *i.e*. search over 𝕆 (3). Looking at the first and fourth images, we see that inclusion of reflections provides a better alignment between the two curves.

**Figure 2:**
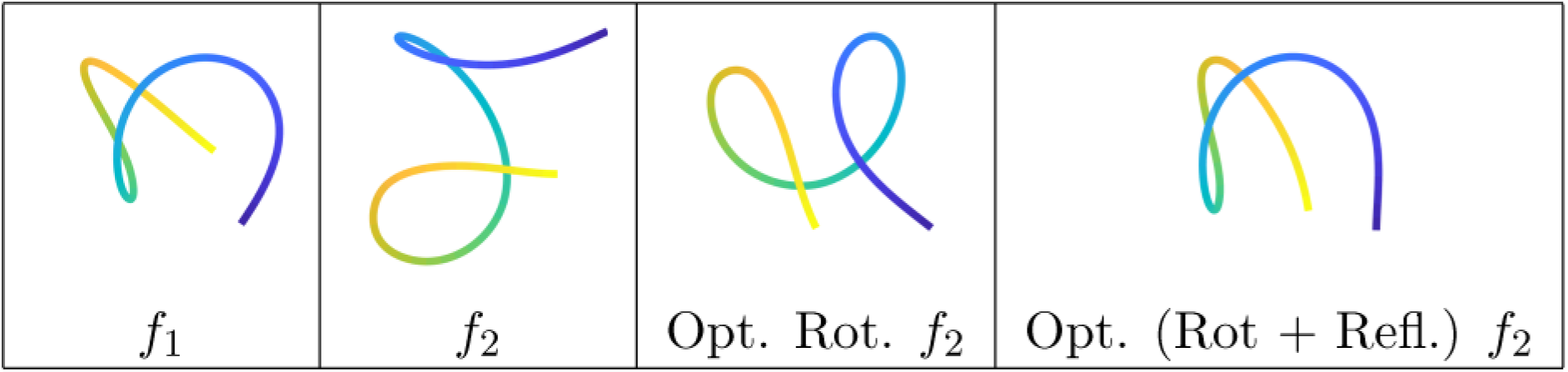
Alignment of *f*_2_ to *f*_1_ using only rotation and using both rotation and reflection.

### 2.2 Clustering of Shapes

The shape metric *d*_*s*_ defined above (Eqn. 1) can be used to compare and cluster curves according to their shapes. Given a set of segments {*f*_*i*_ : [0, 1] → ℝ^3^}, we first compute pairwise shape distances *d*_*s*_(*f*_*i*_, *f*_*j*_) and then use these distances to cluster the segments into groups of similar shapes. This clustering is performed using the `linkage` and `dendrogram` functions in matlab. The number of clusters is determined empirically (manually) by studying the dendrograms. We illustrate this procedure with a simple example in Fig. 3. In this example we cluster 30 spiral curves, nine of which are shown on the right side, using the shape metric *d*_*s*_. These spirals differ in the numbers of loops and pitches, but have the same radii. The image on the left shows the 30 *×* 30 matrix of pairwise distances between them and the dendrogram in the middle shows a perfect clustering of 30 curves into three groups – each group contains curves with the same number of loops.

**Figure 3:**
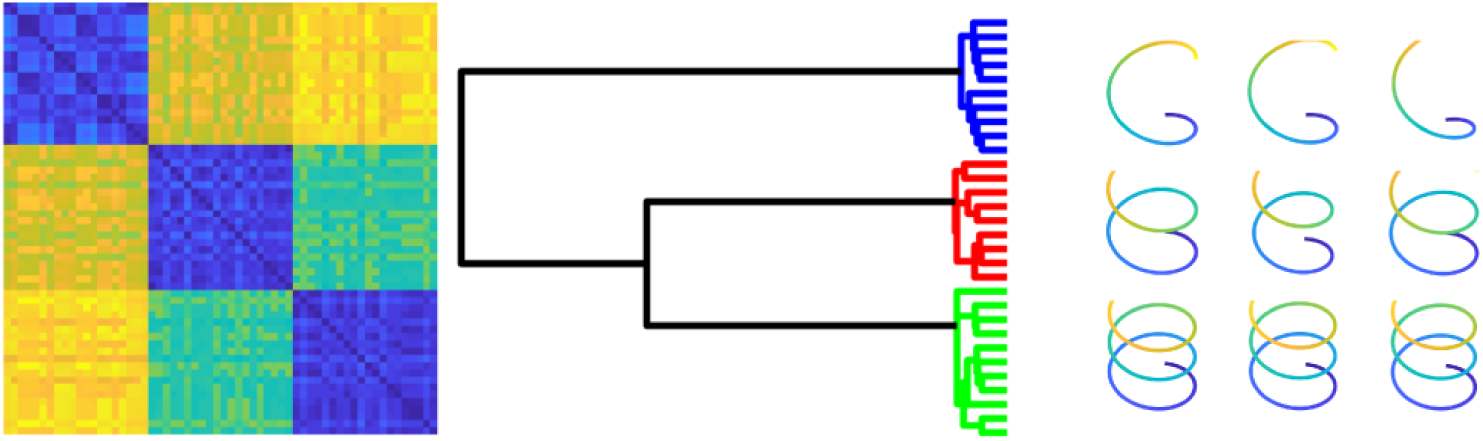
An illustration of clustering 30 curvelets using shape metric *d*_*s*_. The left side shows 30 *×* 30 distance matrix, the middle panel shows their clustering (dendogram), and the right side shows some representative curves from these 30.

### 2.3 Cluster Median

Once we have clustered the curves into homogeneous subgroups, we wish to compute a consensus shape from each cluster. For thus purpose, we can use the notion of a mean or median shape. The median shape, which is a generalization of the geometric median, is an intrinsic statistic that is robust to shape outliers in clusters [5, 20]. A median shape is defined using *d*_*s*_ as follows. Suppose we have a set of *n* parameterized curves {*f*_*i*_}. Their median shape is defined to be the curve

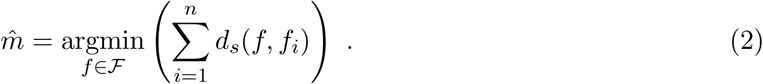

In other words, we find a curve *f* that minimizes the sum of distances to the given curves. Remember that in each distance calculation one has to optimize over rotation/reflections and reparameterizations using SRVF representations. The resulting median is a curve of length one and has no specific orientation or parameterization. In other words, any rotation or re-parameterization of 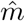 is an equally good solution of Eqn. 2. The algorithm for computing a median shape is a standard one [20], and is not repeated here.

Figure 4 shows an example of this computation. The left side of the figure shows a set of six curves *f*_1_, *f*_2_, …, *f*_6_ – each curve is a spiral with three loops but differs from others in terms of the pitch, radius, and reflection. The right side shows their median shape, which is also a spiral with three loops.

**Figure 4:**
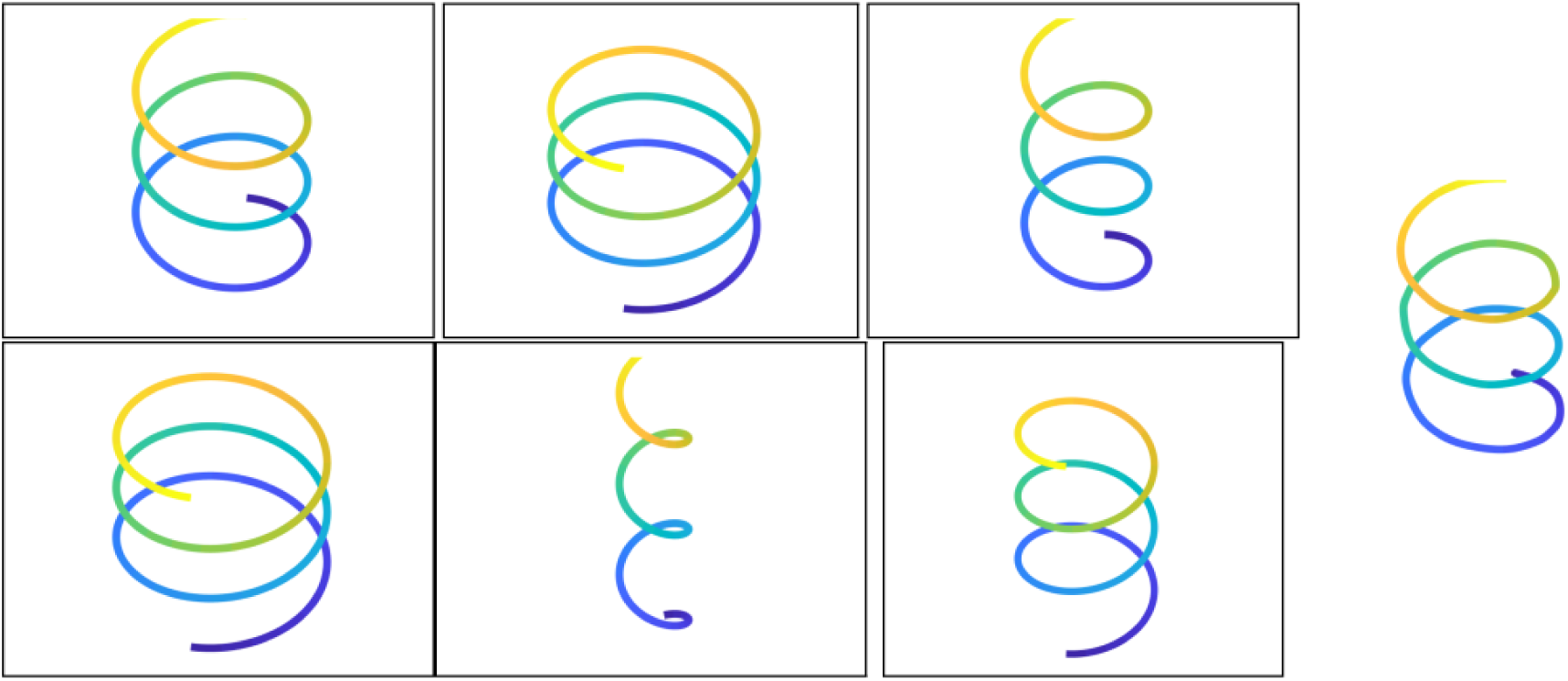
Left side shows six helical curves and right side shows their median shape.

## 3 Constructing Shape Letters

Using shape analysis tools – specifically clustering and averaging – summarized above, we search for representative shapes frequented in segments resulting from chopping training chromosome curves. These shapes then become letters in our CSA. In this section, we describe the step-by-step process of selecting individual shape letters and constructing the full chromosome shape alphabet.

The input to this process is full 3D chromosome curve estimated from Hi-C [2] or imaging data using one or several estimation procedures. Here we use the SIMBA3D package that estimates chromosome shapes from a single cell or ensemble cell Hi-C Data. One can use any other package instead. Assume that for a specific contact dataset, consisting of a number of contact matrices, we have estimated multitudes of chromosome conformations, {*g*_*i*_ ∈ ℱ} using SIMBA3D. We point out that SIMBA3D outputs multiple 3D curves for the same contact matrix, each curve being a local solution to an optimization with random initial condition. Thus, we have hundreds of 3D curves to discover shape letters.

In the experiments presented below, we use a total of 10 contact matrices generated from H9 human stem cells at the 50 KB resolution. These matrices correspond to regions in Chromosomes 1 and 2, and are typically 1000 *×* 1000. We use SIMBA3D to generate 60 3D conformations for each of these contact matrices, 10-20 of which are used for training and the rest are used for testing.

### 3.1 TAD segmentation

Once we have a large set of 3D conformations we segment each one of them into smaller structural units using the concept of TADs. TADs use the structure of a contact matrix to partition it along the main diagonal into disjoint diagonal blocks and then transfers these segmentations to the corresponding curves. Let the *C* be a *p × p* contact matrix and *g* ∈ ℱ be the corresponding 3D curve with *p* points. Let *C*_1_, *C*_2_, …, *C*_*K*_ denote a TAD (block) segmentation of the diagonal of *C* into *K* TADs. Then, chopping *g* into *K* pieces with the same breakpoints as *C* results in *K* curvelets *f* ^(1)^, *f* ^(2)^, …, *f* ^(*K*)^. Several methods, such as TADtree [22] and HiCexplorer [14], have been used to produce TADs. In this paper, we use an insulation score method [19] for creating TADs. Any such method requires the choice of a threshold, analogous to a resolution parameter, which influences the size of the TADs. Due to this, and the dependance of contact matrices on the chosen resolution, we call the eventual CSA *resolution-specific*.

Figure 5 shows an example of this chopping. The left panel shows the image of a contact submatrix with six TAD segmentations overlaid. The right side shows the corresponding six pieces resulting from chopping the larger curve, shown in the middle, at the break points dictated by the matrix segmentation.

**Figure 5:**
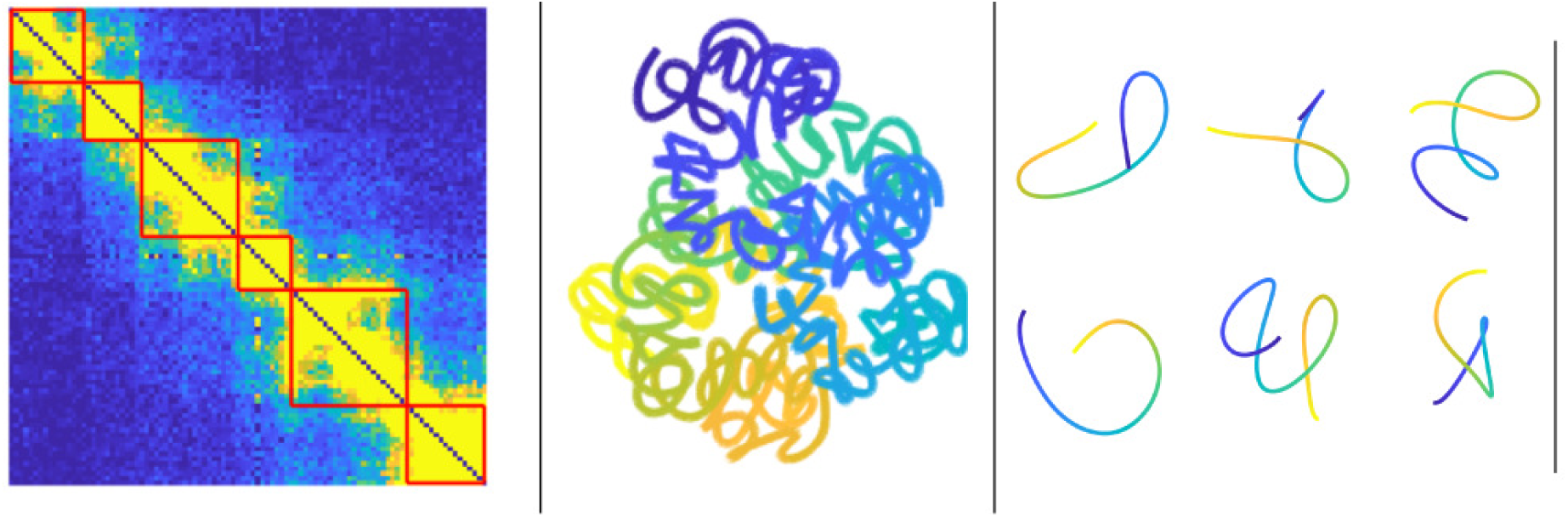
The first six TADs of a contact matrix (left) with the corresponding six pieces (right) obtained by chopping the curve shown in the middle.

### 3.2 Forming a Chromosome Shape Letter

Next, we describe the process of forming chromosome shape letters using the shape toolset covered in Section 2. TAD-based segmentations of all 3D chromosome curves results in a very large pool of segmented curves; call this set 𝒫. Naturally, elements of 𝒫 exhibit a large variability in their shapes. They also differ in sizes, orientations, and parameterizations but these variables do not matter in elastic shape analysis and are easily handled in our approach, as laid out in the previous section. Our goal is to explore 𝒫 and form a subset 𝒜 of distinct, representative shapes that can be used to represent as many observed shapes as possible.

We start by computing pairwise shape distances *d*_*s*_ between all pairs in 𝒫. This is a time consuming process but is performed only once in the learning stage. It results in a large matrix of pairwise distances which is then used to cluster elements of 𝒫. We use the single-linkage hierarchical clustering in matlab to cluster all the curvelets into a number, say *J*, of cluster. The number *J* depends on several factors – distinctness of clusters, desired size of CSA, accuracy of representation, computational resources, and so on.

After clustering elements of 𝒫, we compute a representative shape for each cluster. As mentioned in the last section, we use the notion of a median shape to form this representative shape. A median is chosen because it is relatively robust to the presence of outliers in a cluster. This representative shape then becomes a candidate for being a *shape letter*. In the top part of Fig. 6 we have display four clusters and six curves in each cluster. We compute the median shape of each cluster and display the medians in the bottom row. It is clear that median shapes nicely capture the common structures of curves within the clusters. Also, they seem relatively immune to presence of outlier shapes in the clusters. This process results in a set of shapes 𝒟 = {*δ*_1_, *δ*_2_, …, *δ*_*J*_} that are candidates for CSLs.

**Figure 6:**
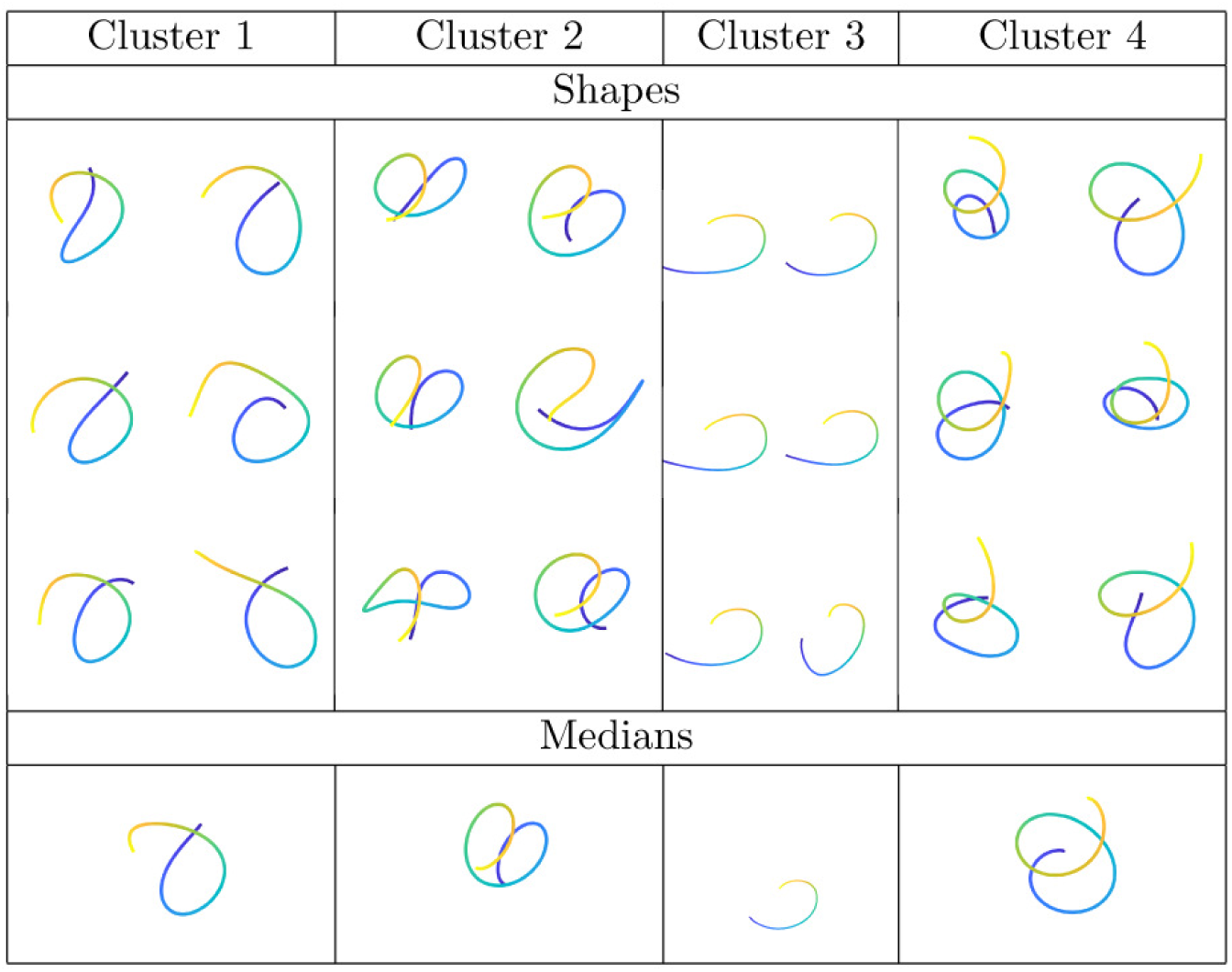
Above: Four clusters with six sample elements each. Below: Shape medians of each of the clusters.

We concede that the results of this stage depend on many choices made – the technique used for TAD segmentation, the chosen number of clusters, the process for clustering, etc. Our hope is that invariances introduced in elastic shape analysis, and the use of robust estimators such as the median, help mitigate, at least to some extent, changes in results due to these choices.

### 3.3 Ordering Shape Letters

It seems convenient to order elements of 𝒟 in certain way that lends to a popular mnemonic usage. One idea is to start with the simplest letter (in the sense of shape complexity) and arrange them in the order of increasing structural complexity. While there are several ways of defining the complexity of a shape, we will use a curvature-based approach. Curvature has several properties that make it useful in this context. It is invariant to translations, rotations, and reflections of a curve, but is not invariant to scale. That is not an issue here since all elements of 𝒟 are of unit length. Also, we remark that the curvature function is not invariant to re-parameterizations. The curvature function of an arbitrarily parameterized curve 3D curve *f* ∈ *F* is given by 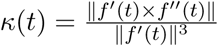.

To quantify structural complexity of curves we calculate its *total absolute curvature*, TAC, given by 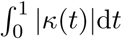. For planar curves, TAC quantifies how far a curve is from being convex [3]. For a general curve, TAC quantifies the amount of bending it goes through. It is important to note that TAC is invariant to not only translation, rotation, reflections but also to re-parameterization. We compute TAC of each of the candidate letter *δ*_*i*_ and reorder them to have increasing complexity. That is, the sorted set 𝒟 = {*δ*_*i*_} has the property that *TAC*(*δ*_*i*_) ≤ *TAC*(*δ*_*i*+1_).

### 3.4 Pruning Shape Alphabet

So far we have an ordered collection of shapes that are potentially our shape letters. The final step is to ensure distinctness of these shapes and remove shapes that are redundant or very similar to other shapes in 𝒟.

We accomplish this by pruning the set 𝒟 sequentially as follows. We shortlist the first letter *δ*_1_ by default; set 𝒜 = {*δ*_1_}. At any iteration, define ℬ to be the complement of 𝒜 in 𝒟. Similar to 𝒟, 𝒜 and ℬ are also ordered sets. Then, we consider the first element of ℬ and move it to 𝒜 if it is at least a certain threshold, say *λ*, from all elements of 𝒜 in terms of the distance *d*_*s*_. Formally, let *δ*_*i*_ be the first element of ℬ and let

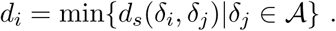

If *δ*_*i*_ satisfies *d*_*i*_ *> λ*, then we move it from ℬ to 𝒜. Otherwise, we just discard it from ℬ. We repeat this procedure until 𝒜 does not change anymore or ℬ becomes empty. In proceeding from left to right in 𝒟, we emphasize a preference for less complex shapes and only accept a new letter in 𝒜 only if it is significantly different from the current letters. This pruning method will remove redundancies in the representation as well as increase efficiency in computation. The process of pruning elements of 𝒟 is summarized in Algorithm 1.

Let 𝒜 denote the resulting set of shape letters. 𝒜 is called the Chromosome Shape Alphabet and elements of 𝒜 are called chromosome shape letters.

#### Algorithm 1 Pruning shape letters from a shape alphabet

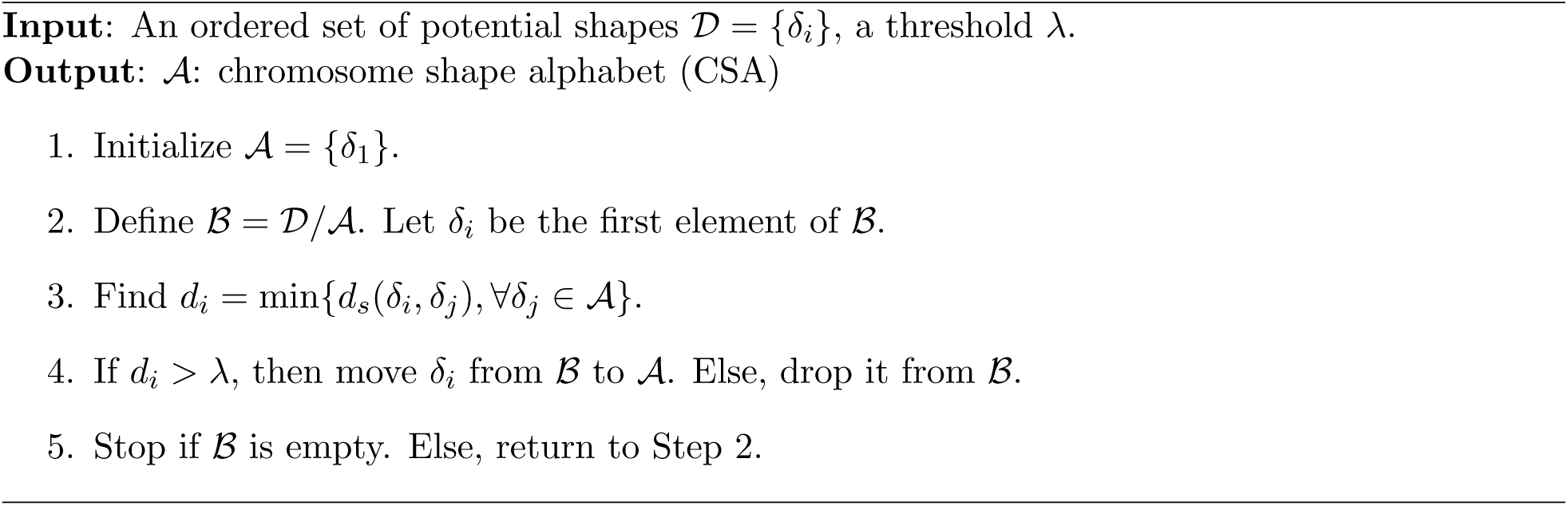

Figure 7 shows the ordered set of 42 CSLs that form our shape alphabet. Note that the first few letters are simple threads that are used to connect more intricate letters that follow in the alphabet. The complexity grows from small bends and twists at the start and reaches three loops plus twists by the end. Even though some letters have the same number of loops, they differ in turns and twists along these loops to result in distinctive shapes.

**Figure 7:**
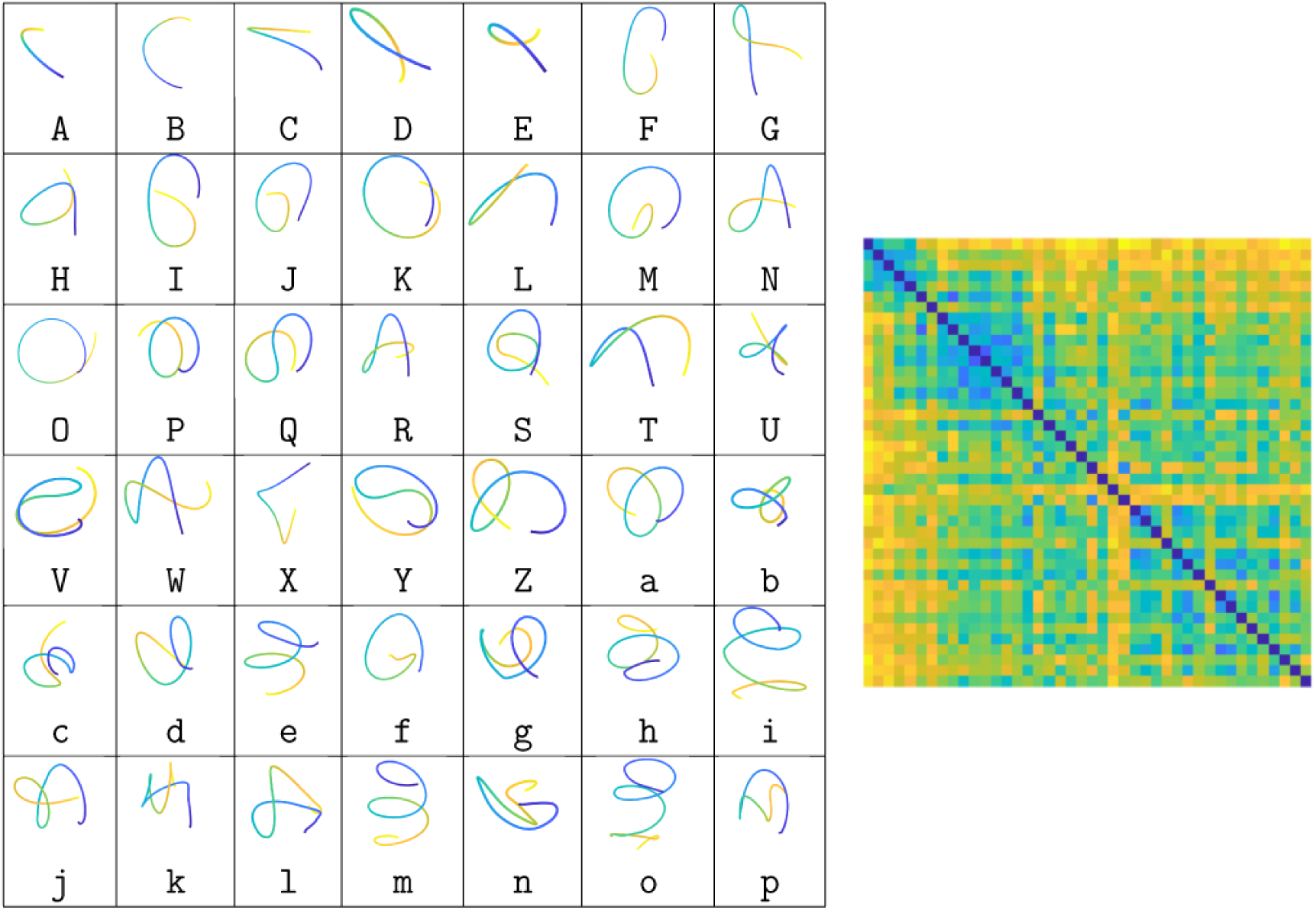
The 42 chromosome shape letters ordered by their shape complexity.

To further study shape differences of these letters we compute pairwise shape distances *d*_*s*_ between these 42 CSLs and display their matrix as an image in the rightmost panel. We observe that the first few letters are quite different from the rest of alphabet. There are a few block structures in the matrix, implying that the letters have some clustering, rather than being “uniformly separated” in the shape space in some way. This alphabet 𝒜 of 42 CSLs is now ready for use in symbolic representation and analysis of chromosomes.

## 4 Alphabetical Representations of Chromosomes

Once we have constructed the shape alphabet, we need to validate it. Since our main goal is to substitute TAD segments, along a full chromosome, by the nearest shape letters, we need to evaluate the errors associated with this substitution. Here we describe the process of forming letter sequences representing chromosomes and reconstructing 3D curves using only the letter shapes. Finally, we discuss the evaluation of these reconstructions. We emphasize that CSA is learnt from the training data while the validation is performed on an independent test data albeit from the same SIMBA3D software.

Suppose *g* ∈ ℱ is a full chromosome 3D curve. a test curve, and let *f* ^(1)^, *f* ^(2)^, …, *f* ^(*K*)^ denote its segmentation obtained using TADs. (Note that we assume that TAD segmentations of these curves are already available for use in the matching.) There are two steps to finding a letter sequence representation of *g*:

- First, for each segment *f* ^(*k*)^ we find the nearest shape letter *δ*_*j*_ ∈ 𝒜. The nearest is defined using the shape distance *d*_*s*_. This results in a string of letters representing that chromosome.
- Second, in order to reconstruct full curves, we need to determine the rotation and the scale of each letter that replaces the corresponding segment. Remember that shape letters represent distinct shapes. There are no rotations, translations, or scales attached to them. One needs to bring these variables back, for each letter usage, in order to put together the full curve again.

We elaborate on these two items next.

### 4.1 Matching Shape Letters

In the first step we find the closest shape letter in our alphabet, for each segment *f* ^(*k*)^, by solving:

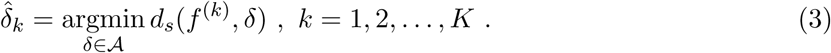

Some examples of these letter matching are shown in Fig. 8. The first row shows six curve segments *f* ^(*k*)^ extracted from a chromosome. The corresponding panels in the middle row show the best matched letters 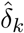 for each of these segments. The matched letters have been rotationally aligned to the top curves to facilitate visual evaluation. The bottom row shows the shape distance *d*_*s*_ between *f* ^(*k*)^ and 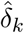. While most letters closely match the observed segments, it is not always the case. For example, the segment *f* ^(5)^ and the matched letter “`g`” have noticeably different shapes. This discrepancy adds to the error in reconstruction. (One can reduce this error by increasing the number of letters in an alphabet as needed.)

**Figure 8:**
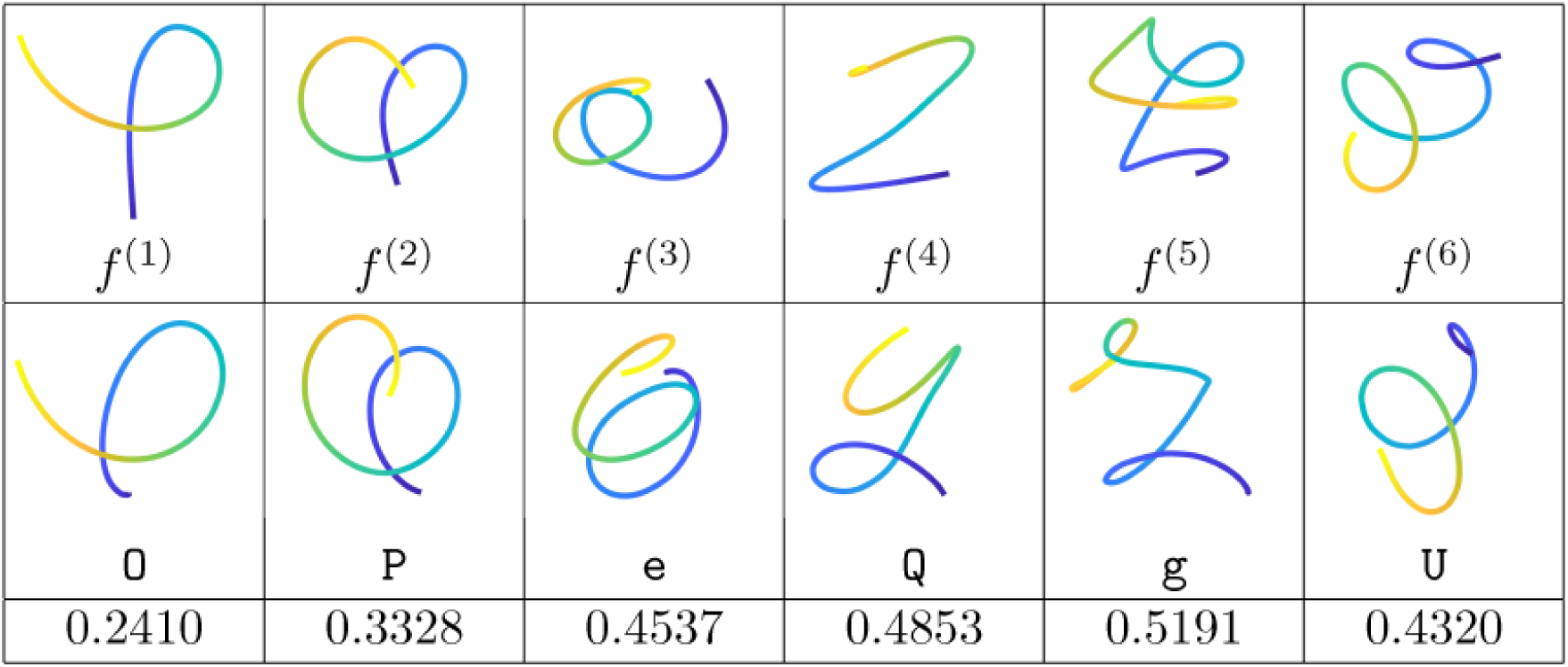
Top: First six segments of a chromosome. Middle: The corresponding closest letters in the shape alphabet. Bottom: the shape distances 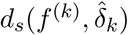.

Using Eqn. 3 for all *k*s, we can obtain a letter sequence representing the whole chromosome 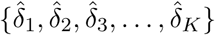. Figure 9 shows five examples of this idea. The top row shows five chromosome structures, and each of the five rows in the bottom shows the corresponding letter sequences for these chromosomes.

**Figure 9:**
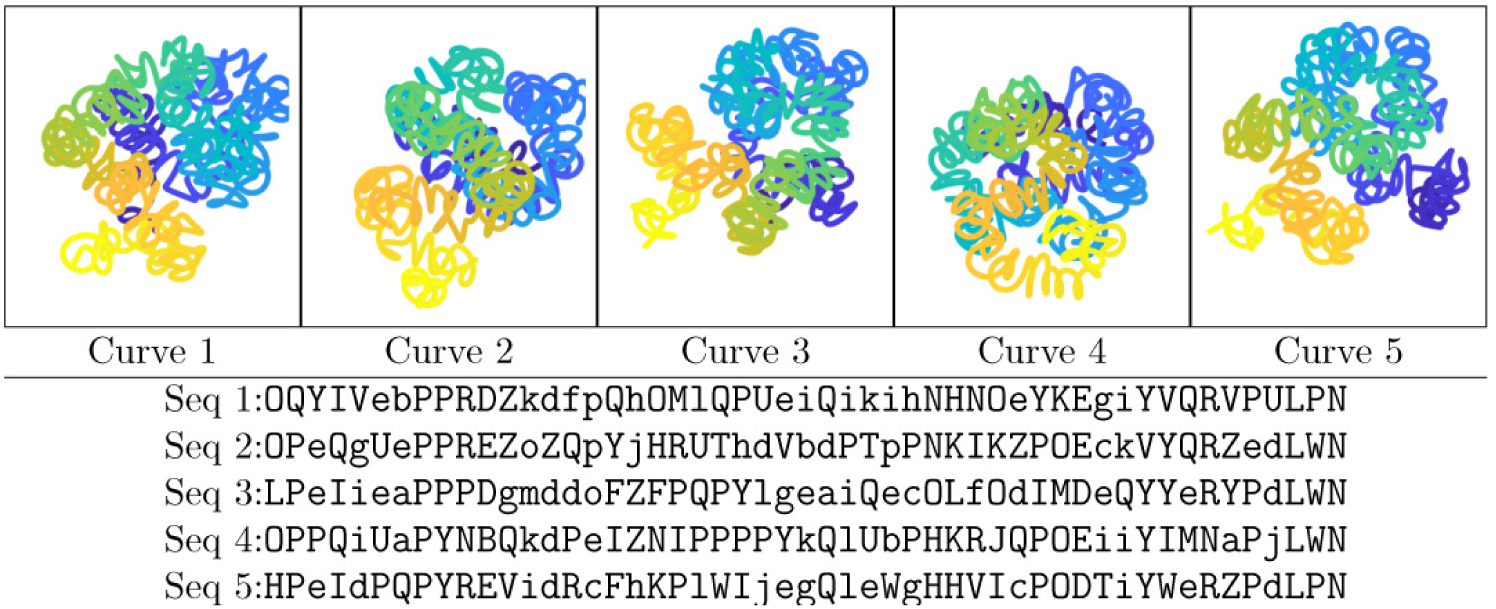
Five chromosome curves that are segmented and represented by CSL sequences shown in the bottom.

### 4.2 CSL-Based Reconstruction: Method 1

The next step is to find optimal transformations (scales and rotations) for each of the letter shapes in order to reconstruct the original curve as faithfully as possible. We start with a simple method that we will term Method 1. For each *k* = 1, …, *K*, we need to find a scale *ρ*_*k*_ ∈ ℝ_+_ and a rotation/reflection *O*_*k*_ ∈ 𝕆 (3) such that the segment 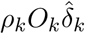 is as similar to *f* ^(*k*)^ as possible. We choose the 𝕃^2^ norm for this comparison and find the optimal transformations using:

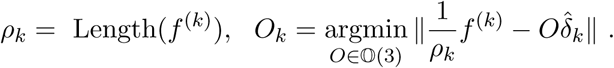

That is, we stretch the shape letter 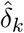 to be the same length as *f* ^(*k*)^ and reorient it with an orthogonal matrix to match the orientation of *f* ^(*k*)^. The reconstructed curve is then simply an ordered end-to-end concatenation of the curve segments 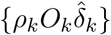, given by:

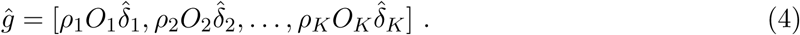

In this comparison, one has to be careful to the number of points being used in represent different segments and letters. It often happens that the segment *f* ^(*k*)^ is discretized with a different number of points than the CSLs. One has to resample either *f* ^(*k*)^ or 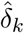 to make sure they have the same number of points, in order to compute the norm. The reconstruction process is summarized in Algorithm 2.

#### Algorithm 2 Reconstruction of a test chromosome

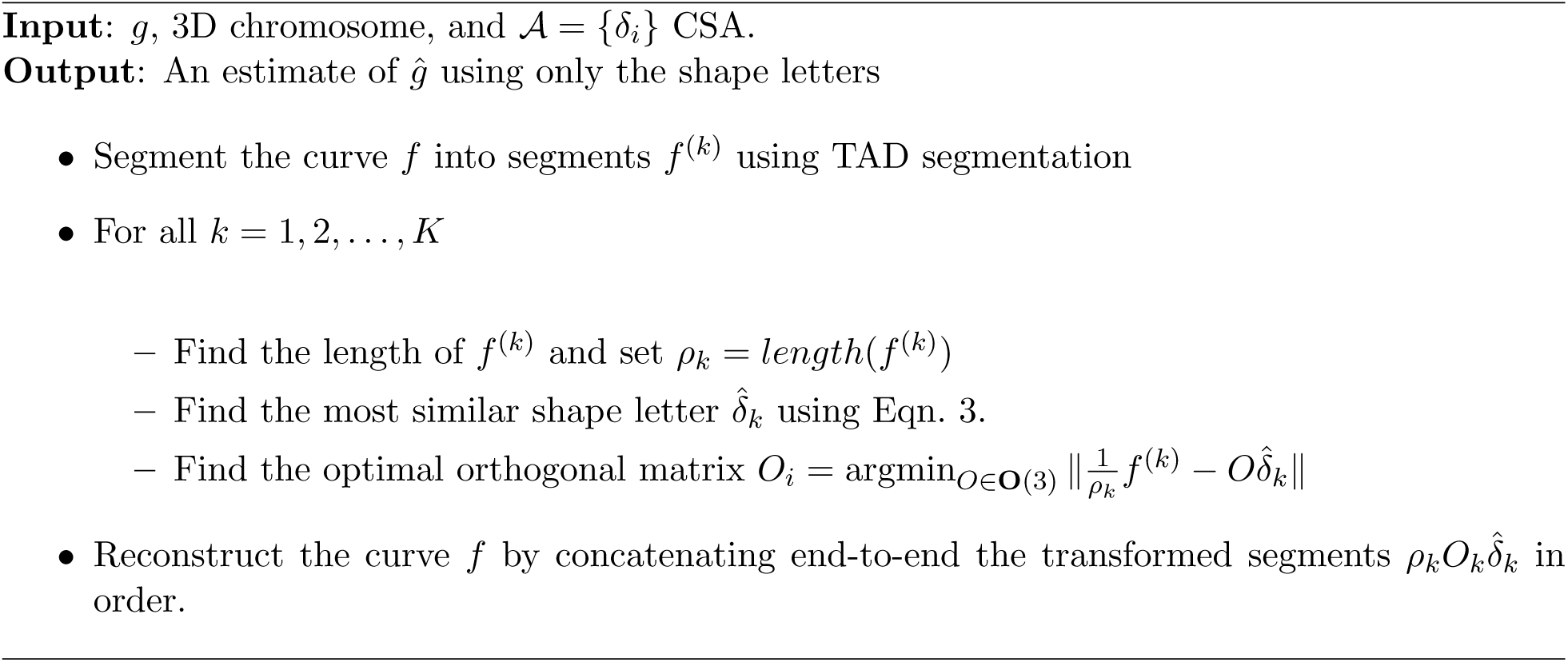

Figure 10 shows some examples of this reconstruction. The top row shows four 3D chromosomes estimated from SIMBA3D package. The corresponding panels in the bottom row show their reconstructions from CSLs using Algorithm 2. Since these shapes are complex, it is difficult to decipher if the reconstruction is successful or not. Some of the reconstructions have similar global shape features to provide some confidence in this reconstruction. In the next section, we discuss a formal evaluation of these reconstructions.

**Figure 10:**
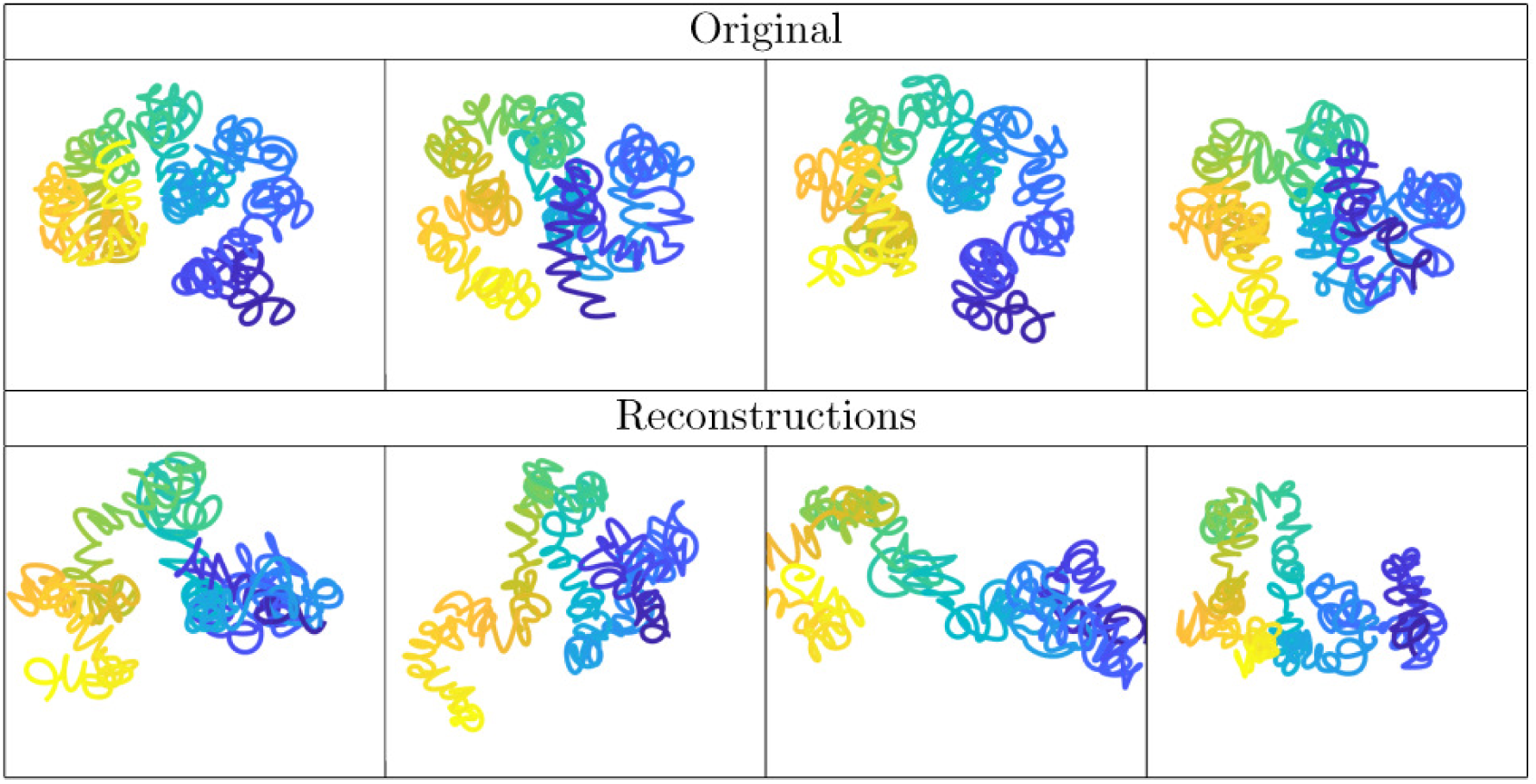
Top row: Four SIMBA3D generated conformations. Bottom row: Corresponding constructions created using only the 42 CSLs and Algorithm 2.

### 4.3 Evaluation

Now that we can reconstruct chromosome curves *g* using *ĝ*, but the question is: How good is this reconstruction? If the reconstructions are found to be consistently good for most chromosomes, then this also validates the choice of shape letters for representing chromosome shape variability. We can evaluate a reconstruction in several ways.

- Shape Metric: For instance, we can compute the shape distance *d*_*s*_ between the original *g* and the reconstruction *ĝ, i.e. d*_*s*_(*g, ĝ*). If the reconstructed full shape is very similar to the original shape then the reconstruction is deemed successful. One possible limitation of this approach is that chromosome structures estimated from contact matrices are often an ensemble of solutions, and not just single distinct shapes. The reconstructed curve can thus differ in shape from the original curve and still be a valid part of this ensemble.
- The other idea is to use the energy/cost that was used in the original estimation of *g* from its contact matrix. For instance, if the original chromosomes were estimated using SIMBA3D, we can use the objective functional employed there to evaluate *ĝ*. In SIMBA3D, this functional is actually the negative Poisson log-likelihood; we shall denote it by *NLL*. If *NLL*[*ĝ*] is similar to *NLL*[*g*], then the reconstruction is considered to be of a good quality.

Taking the second approach, we perform the following experiment. We take 50 curves, originally estimated by SIMBA3D for the same contact matrix, and reconstruct them using a different number of CSLs. The summaries of the negative log-likelihood values or *NLL* are shown in Table 1. We see that as the number of CSLs increases, the *NLL* of the reconstructed curves decreases steadily. Furthermore, if we compare these values with those of the original curves, we see that median *NLL* of reconstructed curves reaches the same order of magnitude as the original curves. This result provides important validation of the CSA approach to representing chromosome shape variability. It also implies that there is a scope for improvement in the reconstruction process.

**Table 1:**
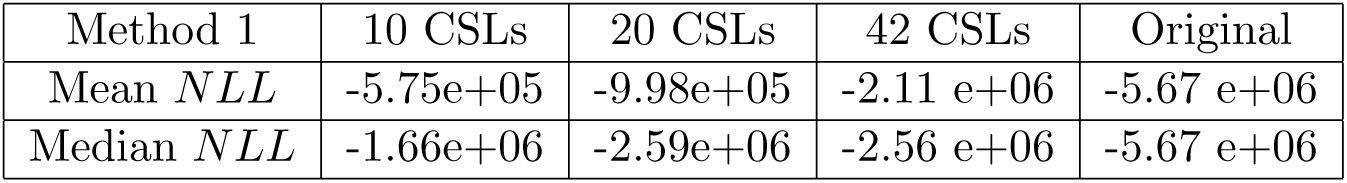
Poisson Negative Log-Likelihoods of *ĝ*, curves reconstructed using only the shape letters, for different number of CSLs used.

**Table 2:** Comparison of *NLL*s of chromosome reconstructions using different methods.

| $NLL$ | Meth. 1 | Meth. 2, $p = 2$ | Meth. 2, $p = 3$ | Original |
| --- | --- | --- | --- | --- |
| Mean | -2.11 e+06 | -5.15 e+06 | **-5.22 e+06** | -5.67 e+06 |
| Median | -2.56 e+06 | -5.17 e+06 | **-5.22 e+06** | -5.67 e+06 |

### 4.4 CSL-Based Reconstruction: Method 2

The method described earlier for computing unknown scales and rotations of CSLs during reconstruction is quite simple. It solves for these transformation for each CSL independently. The other CSLs do not contribute in estimating these transformations for a CSL. Instead of this, one can envision a larger optimization where one solves for all the require transformations, for all the CSLs being used in the reconstruction, in a joint manner. In fact, the objective function for this optimization can be the same *NLL* that was used in SIMBA3D (or any other package being used to generate the original chromosome curves).

We can solve for the unknown transformations jointly by minimizing the objective function

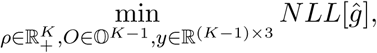

where *NLL* is the negative (Poisson) log-likelihood and *ĝ* is given by:

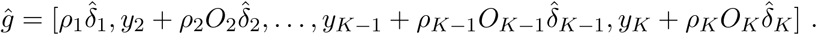

Here *ρ* is a vector of *K* scales, *O* is a set of *K* − 1 rotations/reflections, and *y* is a set of *K* − 1 translations in ℝ^3^. This is a joint optimization problem that adjusts transformations of all CSLs at the same time. Note that one can achieve the same optimal solution without applying a rotation or translation to the first CSL, and thus we leave out *O*_1_ and *y*_1_ from the objective function.

Due to a potential reflection needed for each of the *k* = 2, …, *K* CSLs, executing an exhaustive search over all possible reflection combinations would require 2^*K*−1^ optimizations over 7(*K* − 1) + 1 free parameters. For all but the smallest values of *K*, this approach is generally impractical. There are, however, some greedy approximations that can cut down the computational cost and still provide reasonable solutions. One idea is to break the problem into several subproblems using a sliding window, each involving only *p* neighboring CSLs at a time. For a subproblem, we solve for the optimal scale factor of the first CSL and the full transformation for the remaining *p* − 1 CSLs, by minimizing the *NLL*. This sequential process has the advantage of coordinating transformations across neighbors and still be computationally efficient.

Figure 11 shows examples of reconstructions of chromosomes using the 42-element CSA. We use a sliding window approach in these examples with the window size *p* = 2 in the second row and *p* = 4 in the last row. For a more exhaustive analysis, we repeat the computations of *NLL*, similar to Table 1, but with addition solutions from Method 2. One can see that Method 2 provides substantial improvements in the reconstructions in terms of *NLL* and brings them at par with the original curves.

**Figure 11:**
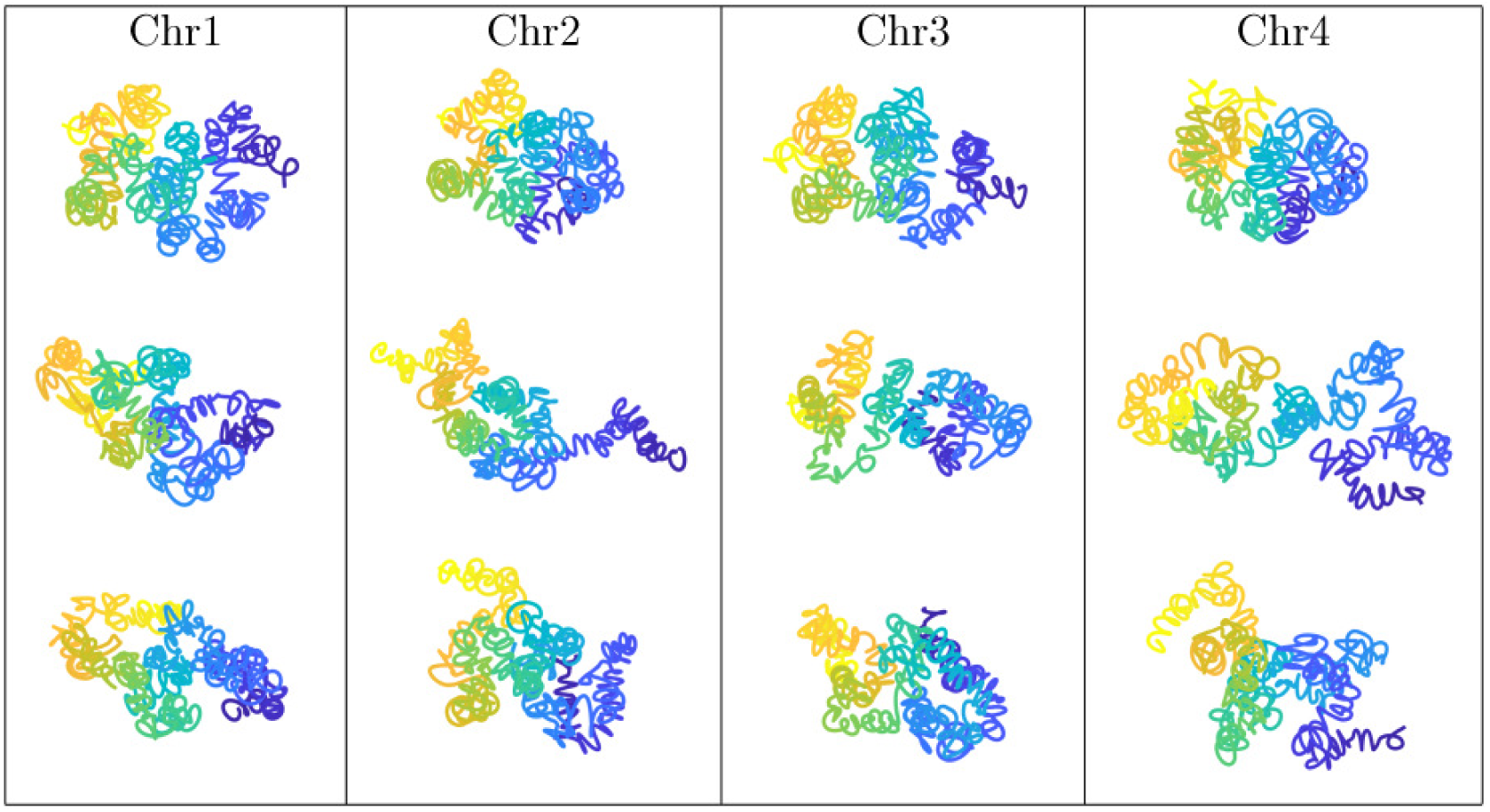
Top row: Four original chromosome conformations from SIMBA3D. Second row: Reconstructions using Method 2. Bottom row: Reconstructions using Method 3.

## 5 Using CSA for Chromosome Structure Analysis

As mentioned at the start, one can use a CSA for abstracting and analyzing shapes of chromosomes in multiple ways. While the main focus on this paper is on the development of a shape alphabet, we present some simple illustrations of their potential usage.

### 5.1 Relative Frequencies of CSLs

As a simple abstraction of chromosome structures into CSLs, we investigate the relative frequencies of different letters in different chromosomes. For this, we take a number of curves generated by SIMBA3D for a contact matrix, represent each of them using strings of CSLs and finally we compute a histogram of how many times each CSL occurs in those sequences. We do not consider the locations of these CSLs but just count their occurrences in the strings.

Figure 12 shows three such histograms associated with three different contact matrices. It is interesting to note both the similarities and differences between these histograms. It is clear from these diagrams that the CSLs do not occur with uniform frequencies in chromosomes. The CSLs in the middle of the alphabet have much higher frequencies. The CSLs with the highest frequency are: `P, Q, I,` and `a`. These shape letters are shown in the bottom panel of Fig. 12.

**Figure 12:**
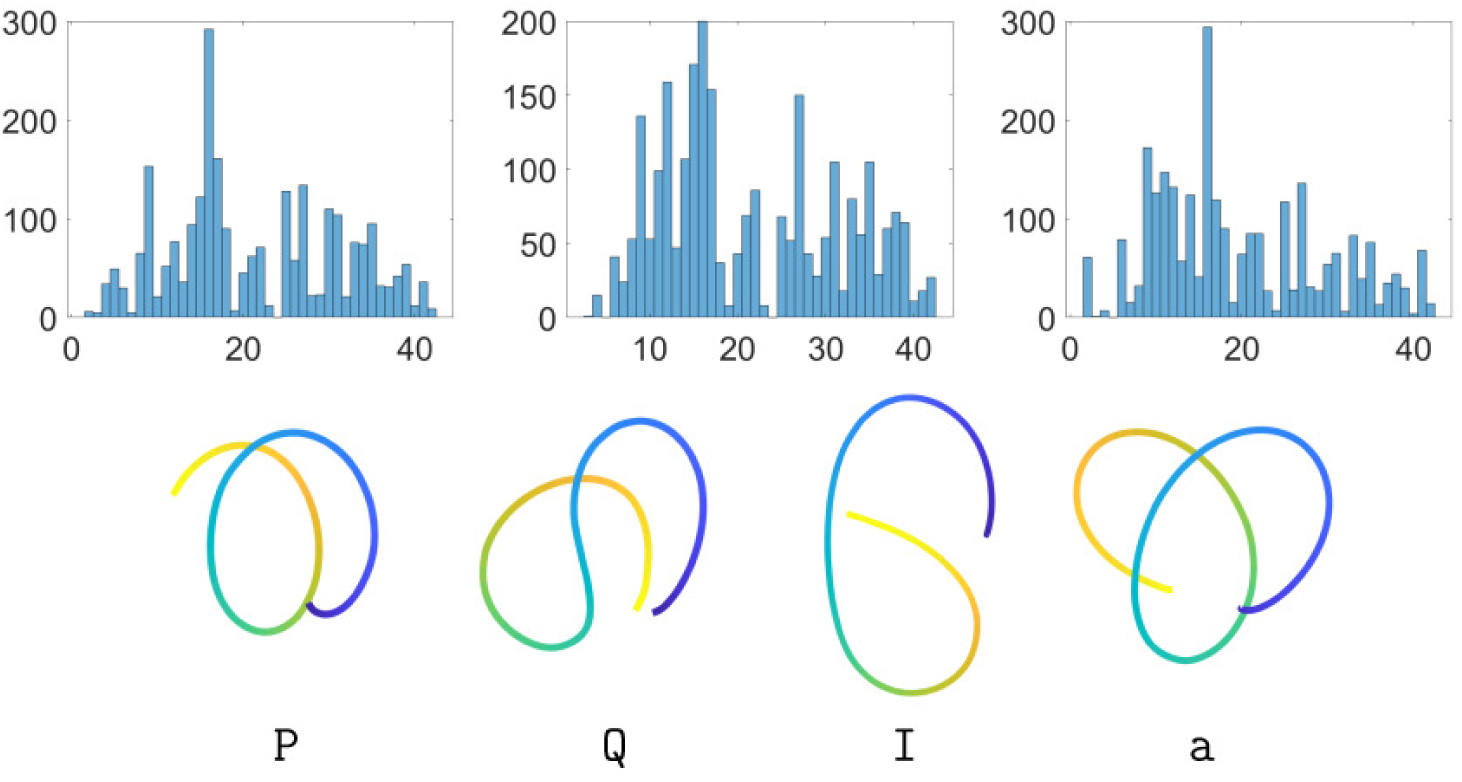
Top: Histograms of appearances of 42 CSLs in SIMBA3D conformations estimated from three different contact matrices. Bottom: Four most common shape letters in these conformations.

### 5.2 Generalized Chromosome Alphabet Logos

Now we have a fully developed framework for representing chromosome shapes with strings of CSLs. It is interesting to “visualize” which symbols frequent the observed shapes and at what locations along the chromosomes. One way to do this is by using sequence logos [18] as follows.

Given a number of symbolic strings, each of length *n*, the sequence logo uses Shannon entropy to measure the uncertainty of having a symbol at a location, say *i*. It computes the height of each letter at that location according to the formula: *Height* = *f*_*b,i*_ *×* (log_2_(*n*) − (*H*_*i*_ + *e*_*n*_)), where 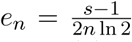, *H*_*i*_ = Σ^*t*^ *f*_*b,i*_ *×* log_2_ *f*_*b,i*_ is the entropy, *f*_*b,i*_ is the relative frequency of shape letter *b* at position *i*, and *s* is the total number of letters. Finally, as a display, it stacks symbols at each location according to their heights. The most frequent symbol is at the top, the next one below it, and so on.

Figure 13 shows a sequence logo associated with 50 SIMBA3D curves generated for a contact matrix. Each curve is segmented into 52 TAD segments, but we display the logo for only the first 15 positions, in order to keep the display readable. In the top panel we show the original sequence logo, where the height and ordering of a letter, at a location, shows its frequency of occurrence in the data at that location. In the bottom part we normalize frequencies at each location so that the heights all add up to one. These sequence logos form a power tool for abstracting and analyzing shape information along chromosome conformations.

**Figure 13:**
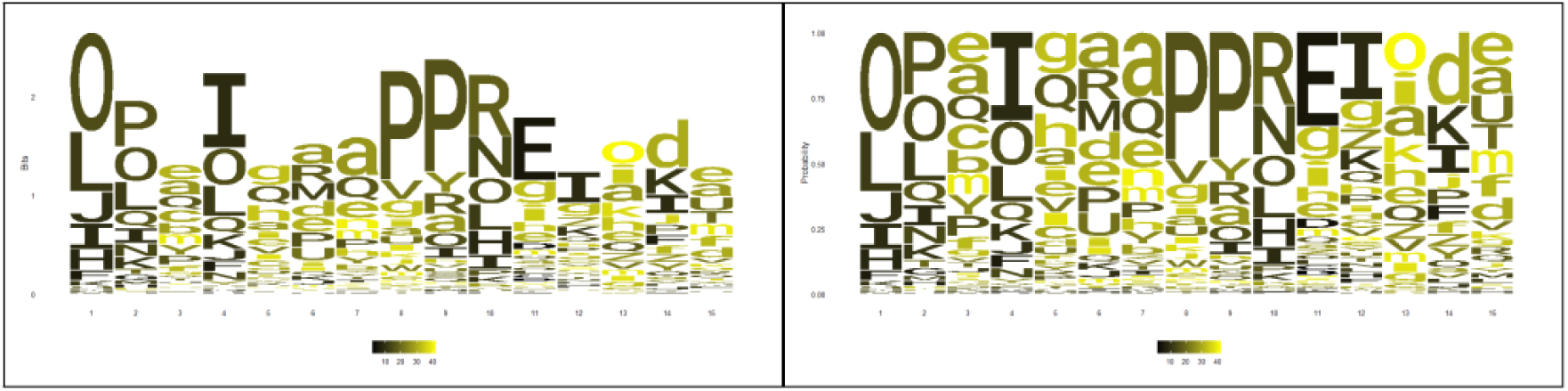
Sequence logos of the first fifteen TADs from a dataset.

## 6 Conclusions

This paper introduces a novel representation of the complex and variable shapes of chromosomes as sequences of shape letters from a shape alphabet. We use shape analysis to lay out a comprehensive step-by-step procedure to construct a shape alphabet from an ensemble of chromosome structures and the corresponding TAD segmentations. As a proof of concept, we demonstrate that this alphabet is sufficient to fully reconstruct chromosomes. Specifically, we show that these reconstructions have negative log-likelihoods that are comparable to the original curves inferred from the Hi-C contact matrix data. We address the problem of structural variability within and across ensembles by using a generalized sequence logo representation of the shape letter frequencies along chromosomes. This manuscript uses ensembles generated as a part of the structural inference process, but the same techniques can be readily applicable to situations where each structure corresponds to an individual cell (e.g. from single cell Hi-C data).

Although we developed our methodology by using chromosome structures inferred from SIMBA3D, any other method that generates chromosome structures can be used. Indeed, it will be important to test how shape alphabets and shape sequence logos will depend on these methods, as well as their dependency on contact map resolution and TAD segmentation. In fact, the alphabet presented here are most likely neither unique nor universal, so it will be important to explore how it varies across cell lines and conditions.

Finally, our novel representation has widespread applicability beyond chromosome structures. For example, we expect it will be applicable to the study of other macromolecules such as unstructured regions of proteins [4] and intrinsically disordered proteins [23], which are gaining a lot of interest in the field.

## Acknolwedgment

This work was supported by the NIH Common Fund Program, grant U01CA200147, as a Transformative Collaborative Project Award (TCPA) to TCPA-2017-NERETTI to NN and AS, NIH R01 GM126558 to AS, NSF CDS&E DMS 1953087 to AS, and NIH/NIA R01AG050582-01A1 to NN.

## References

[1] Bintu B, Mateo LJ, Su JH, Sinnott-Armstrong NA, Parker M, Kinrot S, Yamaya K, Boettiger AN, and Zhuang X. Super-resolution chromatin tracing reveals domains and cooperative interactions in single cells. Science, 362(6413):eaau1783, 2019.

[2] Nynke L Van Berkum, Erez Lieberman-Aiden, Louise Williams, Maxim Imakaev, Andreas Gnirke, Leonid A Mirny, Job Dekker, and Eric S Lander. Hi-c: a method to study the three-dimensional architecture of genomes. JoVE (Journal of Visualized Experiments), (39):e1869, 2010.

[3] Alexander Brook, Alfred M Bruckstein, and Ron Kimmel. On similarity-invariant fairness measures. In International Conference on Scale-Space Theories in Computer Vision, pages 456–467. Springer, 2005.

[4] NE Davey. The functional importance of structure in unstructured protein regions. Curr Opin Struct Biol, 56:155–163, 2019.

[5] P Thomas Fletcher, Suresh Venkatasubramanian, and Sarang Joshi. Robust statistics on riemannian manifolds via the geometric median. In 2008 IEEE Conference on Computer Vision and Pattern Recognition, pages 1–8. IEEE, 2008.

[6] Ming Hu, Ke Deng, Zhaohui Qin, Jesse Dixon, Siddarth Selvaraj, Jennifer Fang, Bing Ren, and Jun S Liu. Bayesian inference of spatial organizations of chromosomes. PLoS computational biology, 9(1):e1002893, 2013.

[7] E. Lieberman-Aiden, N. L. van Berkum, L. Williams, M. Imakaev, T. Ragoczy, A. Telling, Amit, B. R. Lajoie, P. J. Sabo, M. O. Dorschner, R. Sandstrom, B. Bernstein, M. A. Bender, M. Groudine, A. Gnirke, J. Stamatoyannopoulos, L. A. Mirny, E. S. Lander, and Dekker. Comprehensive mapping of long-range interactions reveals folding principles of the human genome. Science, 326(5950):289–293, 2009.

[8] I. Mateo LJ, Murphy SE, Hafner A, Cinquini IS, Walker CA, and Boettiger AN. Visualizing dna folding and rna in embryos at single-cell resolution. Nature, 568(7750):49–54, 2019.

[9] T. Nagano, Y. Lubling, T. J. Stevens, S. Schoenfelder, E. Yaffe, W. Dean, E. D. Laue, A. Tanay, and P. Fraser. Single-cell hi-c reveals cell-to-cell variability in chromosome structure. Nature, 502(7469):59–64, 2013.

[10] T. Nagano, Y. Lubling, C. Vrnai, C. Dudley, W. Leung, Y. Baran, N. Mendelson Cohen, S. Wingett, P. Fraser, and A. Tanay. Cell-cycle dynamics of chromosomal organization at single-cell resolution. Nature, 547(7661):61–67, 2017.

[11] G. Nir and et al. Walking along chromosomes with super-resolution imaging, contact maps, and integrative modeling. PLoS Genet., 14(12):e1007872, 2018.

[12] O. Oluwadare, M. Highsmith, and J. Cheng. An overview of methods for reconstructing 3-d chromosome and genome structures from hi-c data. Biological procedures online, 21(7), 2019.

[13] Jincheol Park and Shili Lin. Impact of data resolution on three-dimensional structure inference methods. BMC bioinformatics, 17(1):70, 2016.

[14] Fidel Ramírez, Vivek Bhardwaj, Laura Arrigoni, Kin Chung Lam, Björn A Grüning, José Villaveces, Bianca Habermann, Asifa Akhtar, and Thomas Manke. High-resolution tads reveal dna sequences underlying genome organization in flies. Nature communications, 9(1):189, 2018.

[15] S. S. Rao, M. H. Huntley, N. C. Durand, E. K. Stamenova, I. D. Bochkov, J. T. Robinson, A. L. Sanborn, I. Machol, A. D. Omer, E. S. Lander, and E. L. Aiden. A 3d map of the human genome at kilobase resolution reveals principles of chromatin looping. Cell, 159(7):1665–1680, 2014.

[16] Michael Rosenthal, Darshan Bryner, Fred Huffer, Shane Evans, Anuj Srivastava, and Nicola Neretti. Bayesian estimation of three-dimensional chromosomal structure from single-cell hi-c data. Journal of Computational Biology, 2019.

[17] Mathieu Rousseau, James Fraser, Maria A Ferraiuolo, Josée Dostie, and Mathieu Blanchette. Three-dimensional modeling of chromatin structure from interaction frequency data using markov chain monte carlo sampling. BMC bioinformatics, 12(1):414, 2011.

[18] Thomas D Schneider and R Michael Stephens. Sequence logos: a new way to display consensus sequences. Nucleic acids research, 18(20):6097–6100, 1990.

[19] Wibke Schwarzer, Nezar Abdennur, Anton Goloborodko, Aleksandra Pekowska, Geoffrey Fudenberg, Yann Loe-Mie, Nuno A Fonseca, Wolfgang Huber, Christian H Haering, Leonid Mirny, et al. Two independent modes of chromatin organization revealed by cohesin removal. Nature, 551(7678):51, 2017.

[20] Anuj Srivastava and Eric P Klassen. Functional and shape data analysis. Springer, 2016.

[21] Q. Szabo, F. Bantignies, and G. Cavalli. Principles of genome folding into topologically associating domains. Science advances, 5(4):eaaw1668, 2019.

[22] Caleb Weinreb and Benjamin J Raphael. Identification of hierarchical chromatin domains. Bioinformatics, 32(11):1601–1609, 2015.

[23] Dyson HJ Wright PE. Intrinsically disordered proteins in cellular signalling and regulation. Nat Rev Mol Cell Biol, 16(1):18–29, 2015.

